# Widespread impact of natural genetic variations in CRISPR/Cas9 outcomes

**DOI:** 10.1101/2023.09.26.559657

**Authors:** Victoria Li, Alicja Tadych, Aaron Wong, Zijun Zhang

**Affiliations:** Harvard University, Cambridge, MA; Lewis-Sigler Institute of Integrative Genomics, Princeton University, Princeton, NJ; Center for Computational Biology, Flatiron Institute, Simons Foundation, New York, NY; Division of Artificial Intelligence in Medicine, Department of Medicine, Cedars-Sinai Medical Center, Los Angeles, CA; Department of Computational Biomedicine, Cedars-Sinai Medical Center, Los Angeles, CA

## Abstract

CRISPR/Cas9 is a genome editing tool widely used in biological research and clinical therapeutics. Natural human genetic variations, through altering the sequence context of CRISPR/Cas9 target regions, can significantly affect its DNA repair outcomes and ultimately lead to different editing efficiencies. However, these effects have not been systematically studied, even as CRISPR/Cas9 is broadly applied to primary cells and patient samples that harbor such genetic diversity. Here, we present comprehensive investigations of natural genetic variations on CRISPR/Cas9 outcomes across the human genome. The utility of our analysis is illustrated in two case studies, on both preclinical discoveries of CD33 knockout in Chimeric Antigen Receptor (CAR)-T cell therapy, and clinical applications of TTR inactivation for treating ATTR amyloidosis. We further expand our analysis to genome scale, population stratified common variants that may lead to gene editing disparity. Our analyses demonstrate pitfalls of failing to account for the widespread genetic variations in Cas9 target selection, and how they can be effectively examined and avoided using our method. To facilitate broad access to our analysis, a web platform CROTONdb is developed, which provides predictions for all possible CRISPR/Cas9 target sites in the coding region, spanning over 5.38 million gRNA targets and 90.82 million estimated variant effects. We anticipate CROTONdb having broad clinical utilities in gene and cellular therapies.

## Introduction

The clustered regularly interspaced short palindromic repeats (CRISPR)/CRISPR-associated protein 9 (Cas9) gene-editing system is highly effective in modifying specific genomic loci. This state-of-the-art gene-editing technology has shown great promise in the treatment of human diseases, including genetic and infectious diseases as well as cancer (1). For instance, β-thalassemia and sickle cell disease (SCD) are the two most common hemoglobinopathies, genetic disorders resulting from mutations that alter the structure of the oxygen-transporting hemoglobin protein. There are ongoing clinical trials (clinicaltrials.gov NCT03745287 and NCT03655678) using CRISPR/Cas9 *ex-vivo BCL11A* erythroid enhancer-disrupted hematopoietic stem cells for the treatment of these hemoglobinopathies, which afflict individuals with diverse ethnic backgrounds (2,3). As of late 2023, the Cas9-based gene therapy to treat SCD has been approved in the UK and the US. Furthermore, CRISPR/Cas9-based T cell editing of human immunodeficiency virus (HIV) co-receptor genes *CCR5* and *CXCR4* could represent an effective cure for acquired immune deficiency syndrome (AIDS), which affects tens of millions of patients worldwide (4,5). CRISPR/Cas9 also has applications in cancer therapeutics, particularly in engineering patient T cells. For example, following the completion of a phase 1 clinical trial (clinicaltrials.gov NCT02793856), deleting the immune inhibitory receptor gene *PDCD1* has been deemed safe and feasible in the treatment of non-small cell lung cancer, the most common type of lung cancer, which is the leading cause of cancer death worldwide (6,7).

CRISPR/Cas9 editing relies on using guide RNAs (gRNAs) to direct a DNA-cleaving endonuclease to precise genomic loci by matching up with an “NGG” protospacer adjacent motif (PAM) sequence. Following a template-free CRISPR/Cas9-induced DNA double stranded break (DSB), cellular repair pathways are activated, which result in nonrandom repair outcomes. However, the exact biological factors affecting these outcomes remains elusive. Studies have shown these outcomes are determined by local DNA sequences, so researchers have turned to machine learning to predict CRISPR/Cas9 editing outcomes (8).

Approaches to this computational biology modeling problem have included neural networks and k-nearest neighbors, multinomial logistic regression, and gradient-boosting decision trees, in the models inDelphi, FORECasT, and SPROUT, respectively (9–11). These approaches rely on manual feature and model engineering, which restrict their effectiveness to limited existing knowledge of CRISPR/Cas9 DSB repair. Automating feature engineering to overcome this limitation motivated the creation of CROTON, a deep residual convolutional neural network (CNN)-based CRISPR/Cas9 editing outcome predictor. CROTON further automated model engineering by leveraging AMBER, a framework built to apply automated machine learning (AutoML)-technique neural architecture search (NAS) to biological CNN design (12). Relative to state-of-the-art inDelphi, FORECasT, and SPROUT, CROTON was more accurate in predicting repair outcomes in a clinically-relevant primary T cell dataset (13).

A major hurdle in therapeutic gene editing is designing appropriate gRNAs for clinical use with efficient on-target editing and effective repair outcomes. In particular, accounting for gRNA off-target effects has been a major research goal, where conventional and interpretable machine learning methods are developed to predict off-target editing sites (14,15). Meanwhile, evaluating editing outcomes following a CRISPR/Cas9-induced DSB has also become increasingly important (11). Furthermore, the prevalence of CRISPR/Cas9 therapeutics-targeted diseases among worldwide populations raises the question of how human genetic variation affects gene-editing outcomes. Notably, single nucleotide variants (SNVs) are the most common type of genetic variation and can be associated with individuals of certain racial or ethnic backgrounds (16).

Currently, most gRNA selection processes are conducted in reference human genomes that do not account for genetic variation, which recent studies have shown may affect therapeutic outcomes. It has been revealed that natural genetic variations can lead to potentially cancerous off-target therapeutic CRISPR/Cas9 editing. These findings have led to the development of online tools like CRISPRme to facilitate variant-aware off-target gRNA assessments (17,18). To further minimize the risks of treatment failure or adverse outcomes in CRISPR/Cas9-based therapeutics related to human genetic diversity, we need to broadly assess how genetic diversity affects on-target CRISPR/Cas9 editing outcomes after a DSB.

CROTON is not only accurate, but also variant-aware. Indeed, it was used in the first analysis of the effect of natural genetic variantions on CRISPR/Cas9 editing outcomes (13). Herein, we present the first systematic computational analysis and evaluation of the effect of human genetic variation on CRISPR/Cas9 gene-editing outcomes. We leverage CROTON to perform computational analyses for therapeutic gRNA design in two case studies –– deleting *CD33* and *TTR* for acute myeloid leukemia (AML) and transthyretin (ATTR) amyloidosis treatment, respectively. To facilitate broad access to our analysis, a web platform CROTONdb is developed, which provides predictions for all possible CRISPR/Cas9 target sites in the coding region. Researchers and clinicians can leverage CROTONdb for the design of effective and broadly applicable CRISPR/Cas9 gRNA targets.

## Materials and Methods

### Genomic data collection and preprocessing

The human reference genome (hg38 release 35) was downloaded from Gencode (https://www.gencodegenes.org/human/) (19). To extract genes and their corresponding coding regions (CDS), we used Gencode comprehensive gene annotation v35 GFF file downloaded from the same website. The python package “gffutils” was employed to analyze the gene annotations and extract corresponding DNA sequences (https://github.com/daler/gffutils). For each CDS in each gene, we extracted all “NGG” sequences from the forward and reverse strands, and extended 30bp in both directions as the candidate Cas9 target region. Within each gene, the candidate Cas9 targets were then sorted based on their smallest genomic coordinates and the corresponding rank was assigned as a numerical PAM ID. For each chromosome, all PAM IDs in this chromosome were compiled into a BED file.

We downloaded the human genomic variations from dbSNP (build 151) from https://ftp.ncbi.nlm.nih.gov/snp/organisms/human_9606_b151_GRCh38p7/ and Gnomad Exome (V2-liftover) from https://gnomad.broadinstitute.org/ in the VCF file format (20,21). We then performed an intersection using BedTools (22) between the genome variation VCF file with the PAM BED file generated from the previous step. The resulting BED file contained the information of PAMs and all overlapping genomic variations, which we used to analyze Cas9-altering variant effect predictions.

### Variant effect prediction

For each pair of a Cas9 target site (with unique PAM IDs) and a genomic variation, we applied CROTON (13), a state-of-the-art deep CNN to predict the Cas9 repair outcomes for the reference and alternative alleles, respectively. Briefly, CROTON is a fully automatically designed model (12) and is trained using a large-scale synthetic library transfected to the K562 cell line (9). When tested in an independent primary T-cell dataset (10), CROTON achieved superior performance, providing the basis for an unbiased variant effect prediction for Cas9-mediated gene therapy. For a given Cas9 target sequence, we used CROTON to predict 1bp insertion probability, and 1bp and 2bp frameshift frequencies. The VEP was defined as the absolute difference between the predicted probabilities of the reference and alternate sequences.

We compiled an indexed, compressed VCF file for each chromosome to store the VEP results. Each row in the VCF file was a variant-PAM combination where we stored the variant location, reference and alternative alleles and sequences, the predicted repair outcomes for the reference alternate sequences. The VCF files were tabix-indexed to enable efficient random access (23).

### Statistical analysis

For each user-queried genomic region or gene, we performed several statistical analyses to characterize the queried region. We analyzed the nucleotide substitution frequencies for all potential Cas9-altering genomic variations and visualized them in a stacked bar plot. We also visualized the frameshift frequency based on the reference sequences and alternate sequences across all PAM sites in the queried region.

To provide the contexts of a user-queried gene or genomic region for Cas9 efficiency and VEP, we pre-computed the genome-wide statistics for frameshift frequencies, variant load, and variant effects. In particular, we computed the cumulative density function (CDF) of Cas9-induced frameshift frequencies across all possible Cas9 targets in the genome, and compared the average predicted frameshift frequency in the user-queried regions to the genome-wide CDF. Similarly, we computed the genome-wide CDF of variant effects across all target sequences. We computed the average per-base-pair variant count and multiplied by the length of the user-queried genomic region to obtain the expected variant count. We fitted a Poisson distribution with its lambda parameter equal to the expected variant count, and used it to derive the percentile for the observed variant count. We then reported all percentiles in the database interface.

### Database implementation

The CROTONdb frontend was implemented in Django, a python-based web framework (https://www.djangoproject.com/). The frontend supports user search by gene name, or by custom genomic region. When searching by gene name, an elastic search engine was implemented to enable fuzzy search and auto completion of gene names. The frontend implementation calls tabix (23) to access the precompiled VCF files in a computational efficient way. Our database is freely available at https://croton.princeton.edu/.

## Results

### Overview of workflow and data analysis

To enable fast and comprehensive investigation of how natural genetic variations could alter CRISPR/Cas9 editing outcomes, we integrated multiple genomic data sources for our computational analysis. We systematically compiled all coding sequences that can be targeted by CRISPR/Cas9 in the human genome, totaling up to 5.38 million gRNA targets. Each of these target sites were assigned with a unique identifier, named PAM ID, based on their genomic coordinates (**Methods**). For existing natural genetic variation, we integrated two popular databases, dbSNP and gnomAD, generating a total of 90.82 million PAM ID/variant pairs (**Figure 1A**).

**Figure 1.**
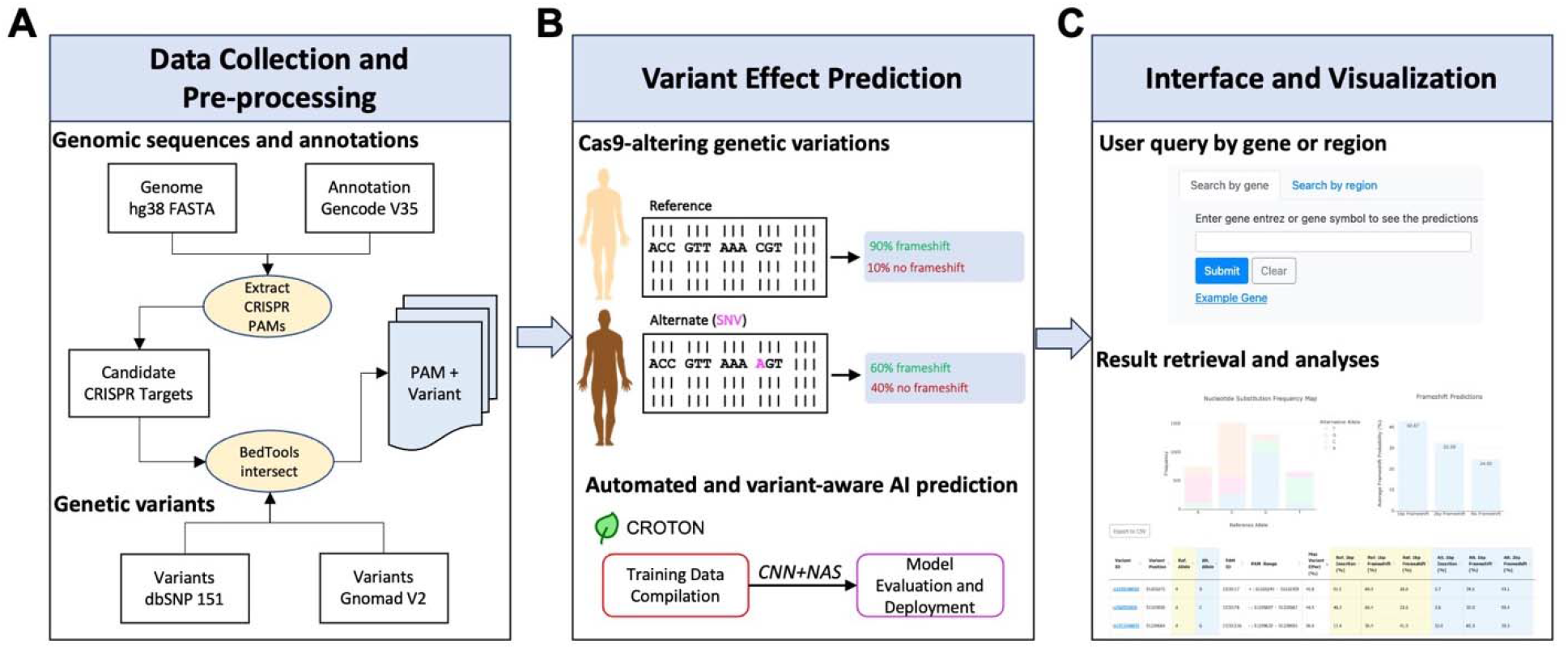
Overview of computational analysis and workflow. **A**) We construct our analyses by comprehensively processing all candidate Cas9 target sites and variant information in the coding region of the human genome. **B**) We leverage CROTON, an automated and variant-aware deep learning model to predict the variant effects for altering Cas9 editing outcomes, which could ultimately lead to differential editing efficiencies. **C**) We then build a web interface, CROTONdb, that provides fast and comprehensive analysis of Cas9-altering genetic variants, enabling users to effectively refine their Cas9 target selection towards efficient and safe genome editing.

The main hypothesis we are testing is whether natural genetic variations, through altering the sequence context of CRISPR/Cas9 target regions, can affect its DNA repair outcomes and ultimately lead to different editing efficiencies (**Figure 1B**). It has been demonstrated previously that the repair outcomes of the DSB induced by CRISPR/Cas9 can be reliably predicted from the local sequence contexts surrounding the target site (8,11). In the hypothetical example, we illustrated that a SNV of reference allele C vs alternative allele A could result in a Cas9-induced frameshift of 90% and 60%, respectively, underscoring a gene knock-out efficiency difference of 30% through the coding region frameshift mechanism. We refer to these variants that can substantially alter the Cas9 outcomes as “*Cas9-altering variants*”. To identify such Cas9-altering variants, we employed CROTON, an automated and variant-aware deep residual convolutional neural network for predicting CRISPR/Cas9 editing outcomes (13). While other machine learning predictors have been developed (9–11,24), we chose CROTON because its convolutional neural architecture does not rely on manually-tuned features, thus capturing the variant effects from a fully data-driven approach. Furthermore, CROTON, designed by state-of-the-art automated deep learning strategy (12), achieved superior performance when compared to conventional machine learning methods.

We illustrate the effect of natural genetic variants in Cas9 editing outcomes with two case studies, covering preclinical discoveries of *CD33* knockout in Chimeric Antigen Receptor (CAR)-T cell therapy, and clinical applications of *TTR* inactivation for treating ATTR amyloidosis. These two case studies highlight the ubiquitous Cas9-altering SNV effects that our analysis identified but may otherwise be missed in existing analysis workflow. We extend this analysis to a genome-scale investigation of CRISPR variant effects across different genetic ancestries. Using this approach, we identify potential population-stratified Cas9-altering variants that should be avoided in Cas9 therapeutic design.

To facilitate the broad access of our analysis to user-specific gRNA design cases, we developed a web tool called CROTONdb (https://croton.princeton.edu/). CROTONdb enables users to query either a gene symbol, or a genomic interval in the human genome (**Figure 1C**). CROTONdb will retrieve all PAM ID/variant pairs within a given gene or an interval in a tabular format, with all predicted Cas9 editing outcomes for the reference and alternative alleles. Analysis for the user’s query will also be performed and visualized automatically. CROTONdb summarizes query predictions by comparing to genome-wide statistics of variant load and effects – users can easily identify and then avoid gene regions where SNVs have large impacts on CRISPR/Cas9 outcomes. This is particularly useful to gRNA designs aimed for broad applicability (e.g., clinically used gRNAs). Furthermore, knockout efficacy of target sequences is predicted and visualized as histograms of frameshift frequencies, enabling prioritization of accurate Cas9 targets. Thus, the CROTONdb web platform can be used as a reference for researchers to avoid SNVs that have a large effect on CRISPR/Cas9 gene editing outcomes.

### Evaluating variant effects in CD33 from preclinical CRISPR/Cas9 discovery

CD33 is a myeloid differentiation antigen that is also expressed on AML blasts and a potential target for cancer immunotherapy. A promising approach to AML treatment is coupling standard bone marrow transplants with CD33-targeted CAR-T Cell therapy. However, since both healthy and malignant myeloid cells are CD33-positive, non-discriminatory therapeutic killing of CD33-expressing cells can compromise immune function and lead to patient death. One approach to addressing this challenge is transplanting CD33-deficient rather than wild type hematopoietic stem and progenitor cells (HSPCs) to patients. These CD33-deficient HSPCs would be spared from CAR T Cell-mediated killing and facilitate functional hematopoiesis (25,26). A recent study has explored this approach, demonstrating that coupling CRISPR/Cas9 knockout (KO) CD33 donor HSPCs with CD33 CAR T Cell therapy is clinically feasible for AML treatment. The gRNAs used in this study were designed with a reference genome and optimized for KO efficiency (25,27).

For this study’s target region-of-interest, we searched for the range “chr19:51225245-51225305” in CROTONdb, then filtering for the study’s PAM ID (i.e. CD33|17) revealed that there existed 18 SNVs within 60 bp of this CRISPR/Cas9 target site. CROTONdb produced a table of corresponding search results with “Export as CSV” functionality that facilitates further investigation of target-associated SNVs, allowing download and easily return to database search results. Using the data table, we analyzed the selected gRNA region relative to the *CD33* gene body and found that the predicted maximum SNV variant effect distribution was more left skewed on the targeted PAM site. While the overall maximum variant effect of *CD33* was concentrated under 10%, our analysis revealed that there are variants with a large effect on CRISPR/Cas9 editing outcomes associated with the study’s target site (**Figure 2A**).

**Figure 2.**
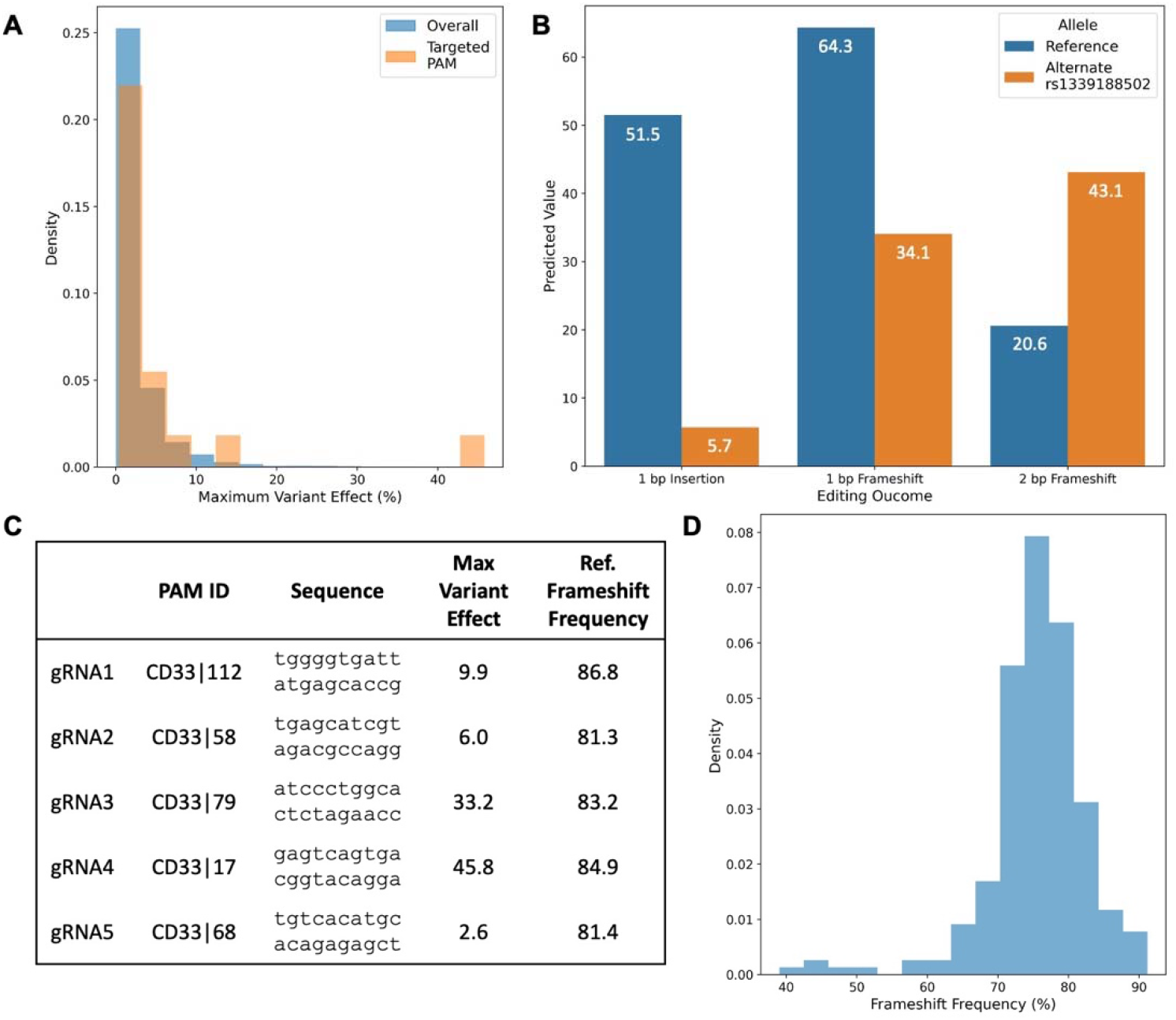
Large variant effects on a reported therapeutic *CD33* CRISPR/Cas9 target site. **(A)** The distribution of the maximum variant effects of SNVs across *CD33* gene and within 60 bp of the proposed gRNA target. **(B)** A comparison of predicted 1 bp insertion, 1 bp frameshift, and 2 bp frameshift frequencies on the reference and alternate genome containing the SNV associated with the highest *CD33* variant effect, rs1339188502. **(C)** The CROTONdb PAM IDs, sequences, and CROTON-predicted maximum variant effect and reference frameshift frequencies for identified efficient CD33 gRNA targets. **(D)** The distribution of predicted frameshift frequencies across the *CD33* gene body.

We sorted the *CD33* results by their max variant effect to identify which target gene regions were associated with the greatest predicted variant effects. Conducting this analysis on *CD33* showed that, in fact, the gRNA chosen by this HSPC KO for AML treatment study was associated with the highest predicted “Max Variant Effect” on *CD33*. More specifically, CROTON predicted that the SNV “rs1339188502,” a missense A to G mutation, decreases the predicted 1 bp insertion probability between the reference and alternate sequences by 45.8%. CROTON’s associated predictions for 1 bp and 2 bp frameshift frequency were also altered substantially by this SNV, decreasing by 30.2% and increasing by 22.5%, respectively (**Figure 2B**).

Especially due to the clinical applications of this work, this observation suggests that a different CD33 gRNA target may be more appropriate for *CD33* inactivation. To select their CD33 target site, the CD33-targeting AML study chose between five gRNAs that had been previously identified as having high CD33 KO efficiency experimentally (25). Evaluating these five gRNAs revealed that they have similar reference frameshift frequencies (i.e., the sum of predicted 1 bp and 2bp reference frameshift frequency predictions) but varying maximum variant effects. Out of these potential targets, the study’s chosen gRNA4 not only had a relatively large reference frameshift frequency of 84.9%, but also the largest maximum variant effect of 45.8% by a wide margin, 12.6% higher than the next largest maximum variant effect. These findings indicate a different gRNA like gRNA1 which had an even higher predicted reference frameshift frequency of 86.8% and a much lower maximum variant effect of 9.9% may be a better choice for therapeutic CRISPR/Cas9 editing for AML treatment (**Figure 2C**). Notably, compared to the distribution of frameshift frequency across the *CD33* gene body, which was concentrated under 80%, the five identified efficient gRNAs had very high predicted reference frameshift frequencies of above 80%. This result substantiates the applicability of using our analysis frameshift frequencies to identify effective gRNAs to target for gene KO (**Figure 2C** and **2D**)

Overall, our machine learning predictions and computation analysis in this case study indicate that focusing on KO efficiency in only reference genomes is ineffective for broad clinical applications. When choosing between gRNAs like the five potential targets for CD33, although researchers found gRNA4 to be most efficacious experimentally, if they had taken variant effects into account, they would have likely chosen a different gRNA target. Tools like CROTONdb, which allow for both sequence and gene-specific variant effect assessments, should be harnessed to facilitate such variant-aware gRNA selection.

### Reprioritizing Cas9 targets in TTR clinical trial to treat ATTR amyloidosis

ATTR amyloidosis is a progressive fatal disease that is increasingly recognized as an important cause of cardiomyopathy and heart failure. This disease is characterized by the accumulation of misfolded transthyretin (TTR) protein in tissues, especially the nerves and heart. Notably, ATTR amyloidosis can be acquired or hereditary –– indeed, 1 of 25 African Americans carry a hereditary *TTR* point mutation that predisposes them to developing this disorder (rs76992529) (28). Current ATTR amyloidosis therapies include reducing or inhibiting TTR protein synthesis by stabilizing tetrameric *TTR* or degrading *TTR* messenger RNA, respectively. These treatments prolong survival but often fail to prevent disease progression and require long-term therapeutic administration.

Recently, researchers conducted a clinical trial targeting ATTR amyloidosis using CRISPR/Cas9-based therapy to inactivate *TTR* (29). They found that following a single infusion of an *in vivo* CRISPR/Cas9 gene-editing therapy targeting patient hepatocyte *TTR*, TTR levels were greatly reduced with mild adverse events (clinicaltrails.gov NCT04601051). In this study, researchers evaluated knockout efficiency, specificity, and on-target editing to select a gRNA targeting *TTR* on chromosome 18 from base pairs (bps) 31592987 to 31593007. CROTON’s predictions on this PAM ID “TTR|24” report a substantial 46% and 30% for 1 bp and 2 bp frameshift frequency, respectively, align with these researcher’s selection for editing efficiency. Indeed, across the *TTR* gene body, the researchers’ targeted site does have relatively high frameshift frequency (**Figure 3A**, highlighted in orange). Notably, there do exist n=19 potential gRNA target sites on *TTR* with a lower maximum variant effect and higher predicted frameshift frequency, including 7 out of a total of 28 on the first two exons of this gene (**Figure 3A**; **Supplementary Table 1**).

**Figure 3.**
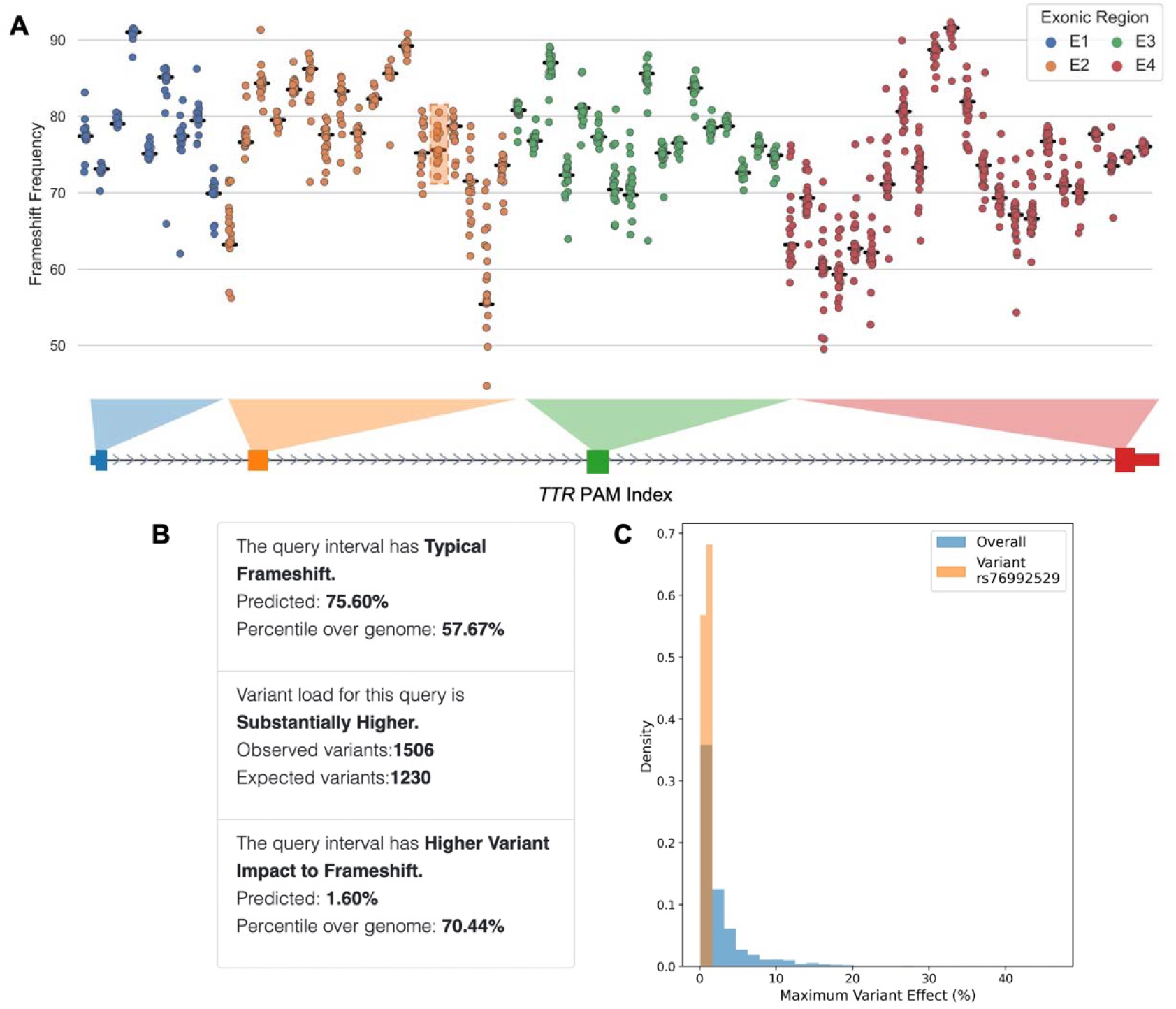
Therapeutic gRNA target analysis across the *TTR* gene body. **(A)** The distribution of predicted frameshift frequency results, the sum of predicted 1 bp and 2 bp frameshift frequency predictions, on n=68 PAM sites across the four coding regions of *TTR*. The orange box denotes the PAM site corresponding to the TTR-targeting ATTR amyloidosis clinical trial, black horizontal lines denote the frameshift frequency prediction for the reference genome, and circles color-coded by exon denote the frameshift frequency predictions for alternate genomes containing SNVs. **(B)** Genome-wide comparisons of the *TTR* target region in terms of frameshift frequency, variant load, and variant impact on frameshift frequency. **(C)** The distribution of the maximum variant effect across *TTR* and corresponding to the rs76992529 SNV, an ancestry-associated genetic SNV that predisposes its carriers to ATTR amyloidosis.

We used the “search by gene” option in CROTONdb to input *TTR* to assess all targetable variant-associated PAM sites on the gene body. Searching by gene also dynamically created graphs that display statistics like predicted frameshift frequencies and variant substitutions that allowed researchers to assess the effectiveness of their gRNA-of-choice against all potential variant-associated target sites in their gene of interest. CROTONdb also outputted statistics comparing the frameshift frequency, variant load, and variant impact to frameshift for the queried regions against the genome-wide statistics (**Figure 3B**).

The genetic links of ATTR amyloidosis further highlight the pertinence of conducting variant-aware gRNA selection in this case. The hereditary *TTR* point mutation rs76992529 that predisposes individuals of African descent to this disease was not linked with high variant effects on potential PAM-targeting sites (**Figure 3C**). We found that that rs76992529 has an average of only 0.92% max variant effects on all its associated Cas9-targeting sites. This impact was much smaller compared to the genetic variations found across the entire *TTR* gene, suggesting that patients will have a comparable editing efficiency regardless of which allele they carry for rs76992529. Nevertheless, variant-aware gRNA selection is still important in this case, as it can help to identify and avoid gRNAs that are likely to be ineffective due to the presence of variants in the target sequence. Attention must be drawn to the variants that are population-stratified or disease predisposing. Our analysis streamlined this process via efficient retrieval and analysis of gRNA frameshift efficiency and all relevant variant information. Based on CROTONdb results, researchers can effectively refine their gRNA selection in order to enable efficient and safe gene therapies with broad applications.

### Mapping the impact of common CRISPR variant effects across diverse genomes

Finally, we substantially expanded our CROTONdb analysis, from existing clinical and preclinical targets to probing the whole-genome scale of Cas9-altering variants. Specifically, we leveraged minor allele frequencies (MAFs) provided in gnomAD to identify population stratification of Cas9-altering variants. Such MAF differences across genetic ancestries poses a particular risk when designing clinical gRNAs and should be preemptively avoided.

We first analyzed the genome-scale distribution of population-stratified Cas9-altering variants. Across the genome, we identified 134,487 genetic variants that were predicted to change the frameshift frequency up to 20%, among which 63,019 variants have been observed in the gnomAD database. We further defined the population stratification as the difference between the highest and lowest MAFs across genetic ancestries larger than 5%, suggesting they are common variants in at least one genetic ancestry, with variable MAFs across populations. Applying this criterion resulted in 777 out of 63,019 population-stratified Cas9-altering variants, corresponding to 1.2% of the total Cas9-altering variants. While population-stratified Cas9-alering variants are observed across all auto chromosomes and the X chromosome (**Figure 4A**), the highest number is from chromosome 1 with 87 such variants, while the lowest number is from chromosome 13 with merely 2 variants.

**Figure 4:**
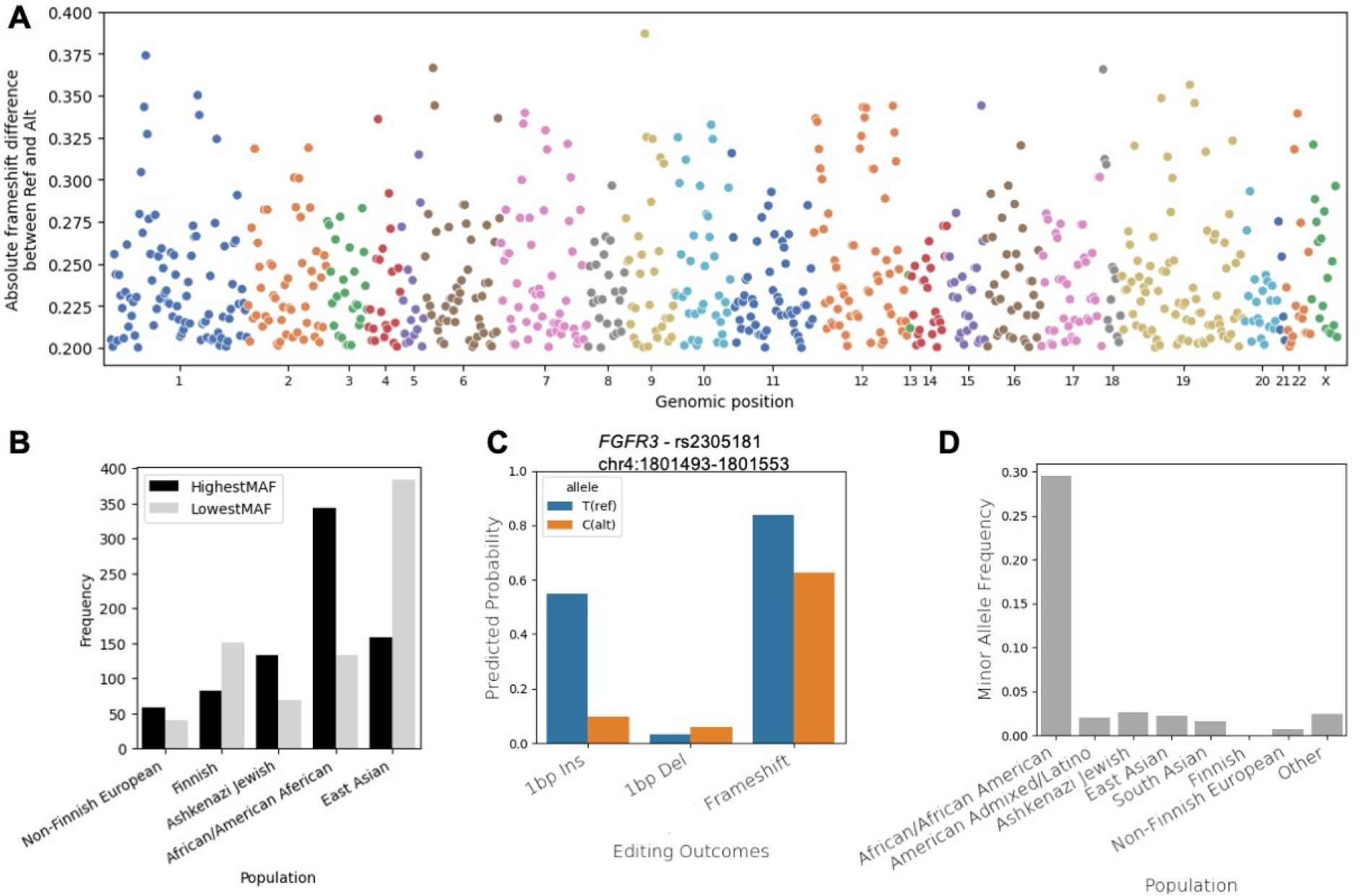
Analysis of genome-wide population-stratified Cas9-altering genetic variants. **A**) Distribution of population-stratified Cas9-altering genetic variants across the whole human genome. **B**) Frequency of different genetic ancestries harboring the highest vs lowest MAFs of Cas9-altering variants. **C**) An example of rs2305181 in *FGFR3*. The predicted CRISPR/Cas9 editing outcomes for sequences with major and minor alleles of the SNV rs2305181. Genomic coordinates of the target region are shown. Ins: insertion; Del: deletion. **D**) minor allele frequencies of rs32305181 across different populations.

A comparison of genetic ancestries with the most disparity MAFs of Cas9-altering variants revealed the African population is potentially vulnerable, if natural genetic variations are not accounted for in Cas9 target design (**Figure 4B**). Across all Cas9-altering variants, African/American African population harbors the greatest number of highest MAF Cas9-altering variants (n=344), indicating they are most likely to carry an alternative allele that alters Cas9 editing outcomes in these loci. Such high MAF Cas9-altering variants are least observed in Non-Finnish and Finnish Europeans, followed by Ashkenazi Jewish and East Asian. In contrast, East Asian population has the greatest number of lowest MAF Cas9-altering variants (n=383), suggesting they may be least susceptible for a substantial proportion (49.2%) among the identified Cas9-altering variants.

As an example, we investigated a candidate Cas9 target in an oncogene gene *FGFR3* that is sensitive to an SNV effect induced by rs2305181 (**Figure 4C**). The minor allele of rs2305181 was predicted to substantially decrease the knockout efficacy from 83.8% to 62.5%. Importantly, rs2305181 has a minor allele frequency of 29.7% and 0.7% in the African and European populations, respectively (**Figure 4D**). If CRISPR/Cas9 is programmed to target this region in primary human cells or in clinical settings, the population stratification of this Cas9-altering variant may lead to health disparities. This demonstrates that this gRNA target should be avoided in the design of widely applicable clinical contexts.

## Discussion

In this study, we developed CROTONdb to search and analyze the potential impacts of natural genetic variations on CRISPR/Cas9 editing outcomes. Through the *CD33* and *TTR* case studies, we demonstrated CROTONdb’s applicability in evaluating PAM sites and discovering additional CRISPR/Cas9 target locations. Notably, CROTONdb’s web features like dynamic graphs and downloadable tables facilitate assessments of the frameshift frequencies and variant-effects of inputted regions-of-interest, facilitating gRNA selection.

With the first CRISPR/Cas9 gene editing therapy to treat cure sickle cell disease being approved in the UK and the US in late 2023, it is expected that other CRISPR-based gene and cellular therapies will follow in the years to come. In conventional medicine and drug treatment, the field of pharmacogenomics has proved valuable in patient care personalization (30,31). The Cas9-altering variants identified in our work are inherently connected with the pharmacogenomics variations that alter drug response in patients. An important distinction to underscore, however, is that patients receiving CRISPR/Cas9 treatments are limited in numbers and will likely remain so for foreseeable future. This restricts the conventional association-based approaches and motivates new computational methods that can preemptively flag potentially problematic target site designs.

CROTONdb is broadly useful to researchers or clinicians applying CRISPR/Cas9 editing to human samples. For basic biological research, CRISPR/Cas9 is often utilized in primary cell lines derived from human donors; in the clinic, therapies that involve CRISPR/Cas9-mediated editing of patient-derived cells have been deemed safe and feasible. These biospecimens all harbor genetic diversity that can lead to differential editing outcomes and eventually genome manipulation or therapeutic efficiencies. Whether in the laboratory or the clinic, CROTONdb has broad utility in ensuring CRISPR/Cas9-based studies and treatments are consistent, accurate, and effective.

## Data Availability

The CROTONdb is publicly available at https://croton.princeton.edu/. The python scripts for constructing the database and performing the analyses can be found in GitHub repo: https://github.com/zhanglab-aim/CROTONdb/tree/main/paper.

