## Supplementary Table 1 for "Widespread impact of natural genetic variations in CRISPR/Cas9 outcomes" (from another version of this preprint: version 1, 2023-09-27)

**Supplementary Table 1: TTR Targets with a Higher Reference Frameshift Frequency and Lower Frameshift Frequency Variant Effect Relative to the ATTR Amyloidosis Clinical Trial Target Site.**

PAM Index 24 (highlighted in yellow) was targeted in the *TTR* clinical trial. This target is associated with a 75.6% predicted reference frameshift frequency and 4.9% predicted absolute frameshift variant effect.

| PAM Index | Variant ID | Variant Position | strand | start | end | Ref. Frameshift | Alt. Frameshift | Exonic Region | Absolute Frameshift Variant Effect |
| --- | --- | --- | --- | --- | --- | --- | --- | --- | --- |
| 4 | rs919662146 | 31591931 | + | 31591904 | 31591964 | 79 | 80.5 | E1 | 1.5 |
| 5 | rs767889884 | 31591939 | - | 31591907 | 31591967 | 91 | 87.7 | E1 | 3.3 |
| 14 | rs528500765 | 31592916 | + | 31592890 | 31592950 | 79.5 | 77.8 | E2 | 1.7 |
| 15 | rs768273993 | 31592924 | - | 31592892 | 31592952 | 83.5 | 87.1 | E2 | 3.6 |
| 19 | rs1383024636 | 31592970 | - | 31592935 | 31592995 | 77.8 | 72.9 | E2 | 4.9 |
| 20 | rs1372138288 | 31592952 | + | 31592935 | 31592995 | 82.3 | 84.3 | E2 | 2 |
| 22 | rs1381337412 | 31592978 | + | 31592956 | 31593016 | 89.2 | 87.2 | E2 | 2 |
| 24 | rs11541796 | 31593011 | + | 31592974 | 31593034 | 75.6 | 80.5 | E2 | 4.9 |
| 29 | rs1397638500 | 31595132 | - | 31595100 | 31595160 | 80.8 | 76.6 | E3 | 4.2 |
| 30 | rs577168116 | 31595134 | + | 31595101 | 31595161 | 76.8 | 79.6 | E3 | 2.8 |
| 33 | rs121918070 | 31595157 | + | 31595127 | 31595187 | 81.1 | 76.2 | E3 | 4.9 |
| 34 | rs121918086 | 31595160 | + | 31595130 | 31595190 | 77.3 | 80.2 | E3 | 2.9 |
| 39 | rs121918071 | 31595209 | + | 31595180 | 31595240 | 76.5 | 74.4 | E3 | 2.1 |
| 40 | rs1804117 | 31595217 | + | 31595184 | 31595244 | 83.7 | 79.9 | E3 | 3.8 |
| 41 | rs1291187509 | 31595231 | + | 31595191 | 31595251 | 78.5 | 80.8 | E3 | 2.3 |
| 42 | rs121918082 | 31595244 | - | 31595208 | 31595268 | 78.7 | 80.1 | E3 | 1.4 |
| 44 | rs121918082 | 31595244 | - | 31595210 | 31595270 | 76.1 | 77.5 | E3 | 1.4 |
| 63 | rs777190076 | 31598640 | + | 31598613 | 31598673 | 76.7 | 72.1 | E4 | 4.6 |
| 66 | rs1333113934 | 31598666 | + | 31598634 | 31598694 | 77.7 | 75.8 | E4 | 1.9 |
| 69 | rs144965179 | 31598662 | - | 31598640 | 31598700 | 76 | 75.1 | E4 | 0.9 |
